# Stool Metabolomics Reveals Catecholamine and Tryptophan-NAD^+^ Pathway Alterations in Autism Spectrum Disorder

**DOI:** 10.64898/2026.02.24.707727

**Authors:** Kevin Liu, Huixi Li, Shaohan Zhang, Muya Xi, Junru Zhu, Jonathan Chen, William Xu, Alex Xie, Alexandros Makriyannis, Jason J. Guo, Xue-Jun Kong

## Abstract

Gut-brain axis dysregulation and microbiome-linked metabolic disturbances have been implicated in autism spectrum disorder (ASD), yet gut-derived neuroactive metabolites remain incompletely characterized. We applied targeted liquid chromatography-tandem mass spectrometry in a cross-sectional case-control study to quantify 18 stool metabolites related to catecholamine synthesis, GABAergic signaling, and tryptophan (Trp) and NAD^+^ metabolism in 59 participants (32 ASD, 27 non-ASD controls). Group differences were evaluated using log_2_ fold change and covariate-adjusted linear models. Random forest classifiers with five-fold cross-validation assessed multivariate discriminative performance, and within-group Spearman correlations examined metabolic network organization. Norepinephrine exhibited the largest elevation in ASD, while dopamine and tetrahydrobiopterin showed nominal group effects consistent with altered BH4-dependent catecholamine metabolism. Among individual metabolites, tetrahydrobiopterin, GABA, and kynurenine were the most informative, and their combination achieved an area under the receiver operating characteristic curve of 0.737 (95% confidence interval, 0.603 to 0.871). Correlation analyses revealed conserved lithocholic acid-deoxycholic acid coupling in both groups. In controls, Trp correlated with kynurenine, whereas ASD showed expanded Trp associations with GABA, norepinephrine, anandamide, and nicotinic acid, consistent with reorganization of Trp-NAD^+^ precursor pathways. These findings identify coordinated, gut-linked metabolic differences in ASD and define a metabolite signature reflecting altered catecholaminergic and Trp-related network structure.

**Importance:** Autism spectrum disorder is currently diagnosed using behavioral assessments, and objective biological measures are limited. The gut microbiome produces many small molecules that can influence brain development and function, but the specific metabolites involved in ASD remain unclear. Using targeted analysis of stool samples, we identified changes in metabolites related to neurotransmitter synthesis and tryptophan metabolism and showed that their relationships form distinct metabolic networks in individuals with ASD. A small panel of metabolites provided moderate ability to distinguish ASD from controls. These results suggest that gut-derived metabolites capture biologically meaningful variation in ASD and may help guide future studies aimed at understanding gut-brain communication and developing noninvasive biomarkers.

## 1. Introduction

Autism spectrum disorder (ASD) is a heterogeneous neurodevelopmental condition characterized by impairments in social communication and restricted, repetitive behaviors. (1) Considerable variability across behavioral, cognitive, and biological domains contributes to the diagnostic complexity of ASD. (2–4) Because diagnosis relies entirely on behavioral assessment (5–7), identifying objective, biologically grounded biomarkers remains a major priority for improving early detection and clinically meaningful subtype stratification.

The gut-brain axis (GBA) has emerged as a promising area for ASD biomarker discovery, as it coordinates the bidirectional communication between the gut and central nervous system and bridges the neural, immune, endocrine, and metabolic signaling pathways that collectively shape the core features of ASD pathophysiology. (8,9) Accumulating evidence has suggested that dysbiotic microbiota are associated with the underlying pathophysiology of ASD through corresponding changes in metabolites. For instance, the restoration of gut microbiota in an idiopathic mouse model of ASD exhibiting autism-like behaviors through human fecal microbiota transplantation has been shown to alleviate social deficit symptoms of ASD, while also normalizing the colon and plasma metabolomes. (10) Parallel evidence involving microbiota transfer therapy in children with ASD has shown persisting improvements in both gastrointestinal (GI) and behavioral symptoms of ASD and a convergence of plasma and fecal metabolome profiles toward those of typically developing children. (11,12) These convergent findings suggest that gut microbial composition and its metabolic activities are functionally coupled with GI and behavioral outcomes in ASD, highlighting the role of microbial metabolites as important mediators of GBA signaling.

Several gut-derived metabolites have been implicated in ASD and provide mechanistic motivation for targeted assays. For example, treatment with *Limosilactobacillus reuteri* increases gut tetrahydrobiopterin (BH4) and restores social behavior in genetic ASD mouse models. (13,14) Microbial influences on catecholamines may alter dopamine (DA) and norepinephrine (NE) balance in ASD. (15) γ-Aminobutyric acid (GABA) has been linked to ASD behavior and microbial composition (16), and altered tryptophan (Trp) metabolism with elevated kynurenine (KYN)-to-Trp ratios has been repeatedly observed in ASD. (17,18) Together, these findings highlight a gut-derived metabolic network linking microbial activity to host neurochemical balance. (19,20)

In this study, we hypothesized that metabolites linked to gut microbiome dysbiosis would exhibit coordinated alterations in ASD and serve as characteristic features of the ASD fecal metabolome. We therefore applied a hypothesis-driven, targeted liquid chromatography-tandem mass spectrometry (LC-MS/MS) approach to quantify gut-derived metabolites previously implicated in GBA signaling. The resulting profiles offer a biochemical window into gut-linked neurochemical processes and support the development of mechanistically grounded biomarkers.

## 2. Methods and Materials

### 2.1. Participants and Ethical Approval

This was an observational, cross-sectional case-control study. The study was conducted in accordance with the ethical principles of the Declaration of Helsinki and approved by the Institutional Review Board of Massachusetts General Hospital (protocol 2022P001749) under a formal collaboration agreement with Northeastern University (agreement 2023A064744). Parents or legal guardians of all participants provided written informed consent, and child assent was obtained when developmentally appropriate.

Participants with ASD were recruited from Massachusetts General Hospital (Charlestown, Massachusetts, USA) based on the following inclusion criteria: (1) male or female participants aged 2.5-22 years; (2) confirmed clinical diagnosis of ASD; (3) no recent use of antibiotics, probiotics, or oxytocin within the previous 4 weeks; (4) stable medication regimen for at least 4 weeks prior to enrollment; and (5) willingness to provide stool samples and participate in interviews and study procedures. The exclusion criteria included the following: (1) pregnancy (before or during the study); (2) comorbid neurological or psychiatric disorders such as bipolar disorder or history of substance use disorder, active cardiovascular disease not controlled by medication, or current use of psychotropic medications; and (3) treatment with oxytocin or probiotics within 4 weeks prior to sample collection.

Non-ASD controls (NAC) were recruited from the community and research registries and the inclusion criteria were: (1) male or female participants aged 2.5-22 years; (2) no diagnosis of ASD or other developmental disorder; (3) no recent use of antibiotics, probiotics, or oxytocin within the previous 4 weeks; (4) stable medication regimen for at least 4 weeks; and (5) willingness to provide stool samples and participate in study procedures. Exclusion criteria mirrored those for the ASD group, including pregnancy, comorbid neurological or psychiatric disorders, active uncontrolled cardiovascular disease, and recent treatment with oxytocin or probiotics.

ASD diagnosis was defined based on a confirmed clinical diagnosis documented in the medical record. Non-ASD controls were defined as individuals without a diagnosis of ASD or other developmental disorder. Age and sex were obtained from parent report and medical records. No interventions were administered as part of this study.

### 2.2. Chemicals and Reagents

HPLC-grade methanol, acetonitrile, water, and formic acid (98%) were purchased from Fisher Scientific (Hampton, NH, USA). Internal standards, including deoxycholic acid-d4, tryptophan-d5, serotonin-d4, and dopamine-d2, were obtained from C/D/N Isotopes (Vaughan, ON, Canada). All 18 chemical metabolites of interest were acquired from Thermo Fisher or MilliporeSigma, except anandamide and 2-arachidonoylglycerol, which were prepared internally at the Center for Drug Discovery, Northeastern University. These compounds were first used to generate standard calibration curves on the mass spectrometer and were then prepared as quality-control (QC) mixtures to monitor instrument stability and reproducibility while analyzing the study samples.

### 2.3. Sample Collection and Preparation

Fresh fecal samples were collected from participants in fecal collection and storage tubes containing ethanol (95%) (OMNImet•GUT, DNA Genotek, Canada) and stored at -80 °C until analysis. For metabolomic profiling, samples were thawed at 4 °C and prepared in triplicate. Approximately 30 mg of fecal material was mixed with 400 μL of ice-cold methanol, homogenized at 25 Hz for 5 min, and centrifuged at 10000 × g for 10 min at 4 °C. The supernatant was transferred to a clean tube and dried using a Labconco CentiVap concentrator (Labconco, Kansas City, MO, USA) for 40 min at 35 °C. The dried residue was re-suspended in 150 μL methanol:water (1:1, v/v), spiked with 10 μL of the internal standard mixture, and centrifuged again for 3 min at 10000 × g (4 °C). Processed samples were stored at -80 °C until LC-MS/MS analysis.

### 2.4. LC-MS/MS Analysis

LC-MS/MS analyses were performed at Northeastern University (Boston, Massachusetts, USA). Chromatographic separation was performed on a Vanquish UHPLC system coupled to a TSQ Altis™ triple-quadrupole mass spectrometer (Thermo Fisher Scientific, San Jose, CA, USA). The injection volume for all samples was 20 μL.

#### 2.4.1. Chromatographic Conditions

Metabolites were separated using a Kinetex® F5 column (3.0 × 150 mm, 2.6 μm) with a SecurityGuard™ ULTRA UHPLC F5 guard column, maintained at 30 °C. The mobile phase consisted of 0.1% formic acid in water (A) and 0.1% formic acid in acetonitrile (B). The gradient elution program was run at 0.7 mL/min as follows: 0-2 min, 10% B; 2-10 min, 5-95% B; 10-12 min, 95% B; 12-13.5 min, 95-10% B; and 13.5-15 min, 10% B for re-equilibration.

#### 2.4.2. Mass Spectrometry Conditions

The mass spectrometer was operated in both positive and negative electrospray ionization (ESI) modes using Thermo Fisher’s Advanced Active Ion Management (AIM) polarity-switching technology. Source parameters were optimized as follows: positive ion voltage 4500 V; negative ion voltage 3000 V; sheath gas 60 (arb); auxiliary gas 7 (arb); sweep gas 2 (arb); ion-transfer tube temperature 380 °C; vaporizer temperature 350 °C. Collision energies and fragment ions were optimized for each metabolite using authentic standards or predicted from molecular structures (Table S1).

#### 2.4.3. Quality Control

A quality-control (QC) dilution series was prepared from a mixture of all metabolites of interest at four concentrations to span the expected dynamic range: blank (solvent only), low QC (0.025 μM), medium QC (0.6 μM), and high QC (7.5 μM). QC samples were injected after every 40 study samples to monitor system stability, reproducibility, and analytical performance. Blank samples were used to assess background contamination and matrix effects.

### 2.5. Data Processing

All quantified metabolite concentration data were processed using *MetaboAnalyst* (version 6.0) (21) . For each biological sample, technical replicate intensities (n = 3) were first smoothed using kernel density estimation with a Gaussian kernel (bandwidth = 0.5) to reduce technical variability. These smoothed values were then averaged using the arithmetic mean. To ensure data quality, a filtering step was applied such that, within each sample, any metabolite feature with more than 33% missing values and a coefficient of variation (CV) greater than 1 across replicates was assigned a value of zero. After averaging, normalized data were obtained using a sequential pipeline comprising median normalization to account for inter-sample variability, log10 transformation to reduce right skew and compress the dynamic range, and Pareto scaling to emphasize moderate-intensity changes while retaining biological interpretability.

### 2.6. Statistical Analysis

All statistical analyses were performed in *R* (version 4.4.2). Machine-learning workflows were implemented using the *tidymodels* framework (version 1.3.0), with random forest models fitted using the *ranger* engine (version 0.17.0). Classification performance metrics were computed using *yardstick* (version 1.3.2), and receiver operating characteristic analyses were conducted using *pROC* (version 1.19.0.1). Statistical testing was performed using *rstatix* (version 0.7.2). Pairwise correlations were computed using the *Hmisc* package (version 5.2.3). Data visualization was conducted using *tidyverse* packages. Unless otherwise specified, default parameters were used.

Age was analyzed as a continuous variable, sex was analyzed as a binary categorical variable, and diagnostic group was coded as ASD versus non-ASD control. Factors were coded such that NAC and Female served as reference levels. For descriptive summaries, measurements below the limit of detection (LOD) were excluded. For inferential and machine-learning analyses, below-LOD values were retained as zeros in the normalized data matrix to preserve group structure.

#### 2.6.1. Differential Abundance

Metabolite concentrations were median-normalized, log10-transformed, and Pareto-scaled as described above. Scaling was applied primarily for multivariate modeling to prevent high-abundance metabolites from dominating model training. Group differences were assessed using Wilcoxon rank-sum tests and linear models adjusted for age and sex. Log2 fold changes were derived from differences in log10-transformed means and converted to log2 scale by multiplying by log2(10). P-values were corrected using the Benjamini-Hochberg false discovery rate (FDR) method with significance defined as q < 0.05.

#### 2.6.2. Correlation Analysis

Within-group associations were assessed using Spearman rank correlation coefficients. FDR correction was applied separately within each group across all tested metabolite pairs.

#### 2.6.3. Machine Learning

Random forest classifiers were constructed using 500 trees with default mtry and minimum node size parameters. No hyperparameter tuning was performed due to limited sample size. Five-fold cross-validation stratified by diagnostic group was used to evaluate model performance. A fixed random seed (123) ensured reproducibility.

Model performance was assessed using AUC, accuracy, sensitivity, specificity, precision, recall, F1-score, and Brier score. ROC curves were generated using cross-validated predicted probabilities pooled across folds. AUC confidence intervals were estimated using 2,000 bootstrap resamples. ASD was defined as the positive class. Class weights were not applied. Calibration was assessed using LOESS-smoothed curves comparing predicted probabilities to observed outcomes.

### 2.7. Reproducible Research and Data Availability

All statistical analyses and data visualization are reproducible using the source code available at https://github.com/kevinliu-bmb/targeted-asd-stool-metabolomics.

Raw LC-MS/MS vendor files, processed metabolite abundance matrices, and anonymized participant-level metadata are not publicly available due to human-subjects protections and institutional restrictions. Access to these data may be considered upon reasonable request and subject to Institutional Review Board approval and execution of a data use agreement. Requests should be directed to the corresponding author.

### 2.8. STORMS Checklist

The STORMS checklist for this publication can be found at https://github.com/kevinliu-bmb/targeted-asd-stool-metabolomics/blob/main/STORMS_checklist.xlsx.

## 3. Results

The present study included a total of 59 participants, of whom 32 individuals aged 6.86 ± 4.66 [2.72, 19.81] years were diagnosed with ASD, and 27 individuals aged 8.57 ± 6.11 [2.43, 21.56] years were NAC. The ASD group consisted of 24 males and eight females, whereas the NAC group comprised 10 males and 17 females. The age distribution of participants in each group is shown in Figure S1. Measured fecal metabolites are summarized in Table 1, and their groupwise distributions are shown in Figure 1. Notably, the quantification for LCA, NE, and UA have yielded measurements that are below the LOD consistently among a number of sample triplicates for certain participant samples, and these details are also summarized in Table 1.

**Table 1.**
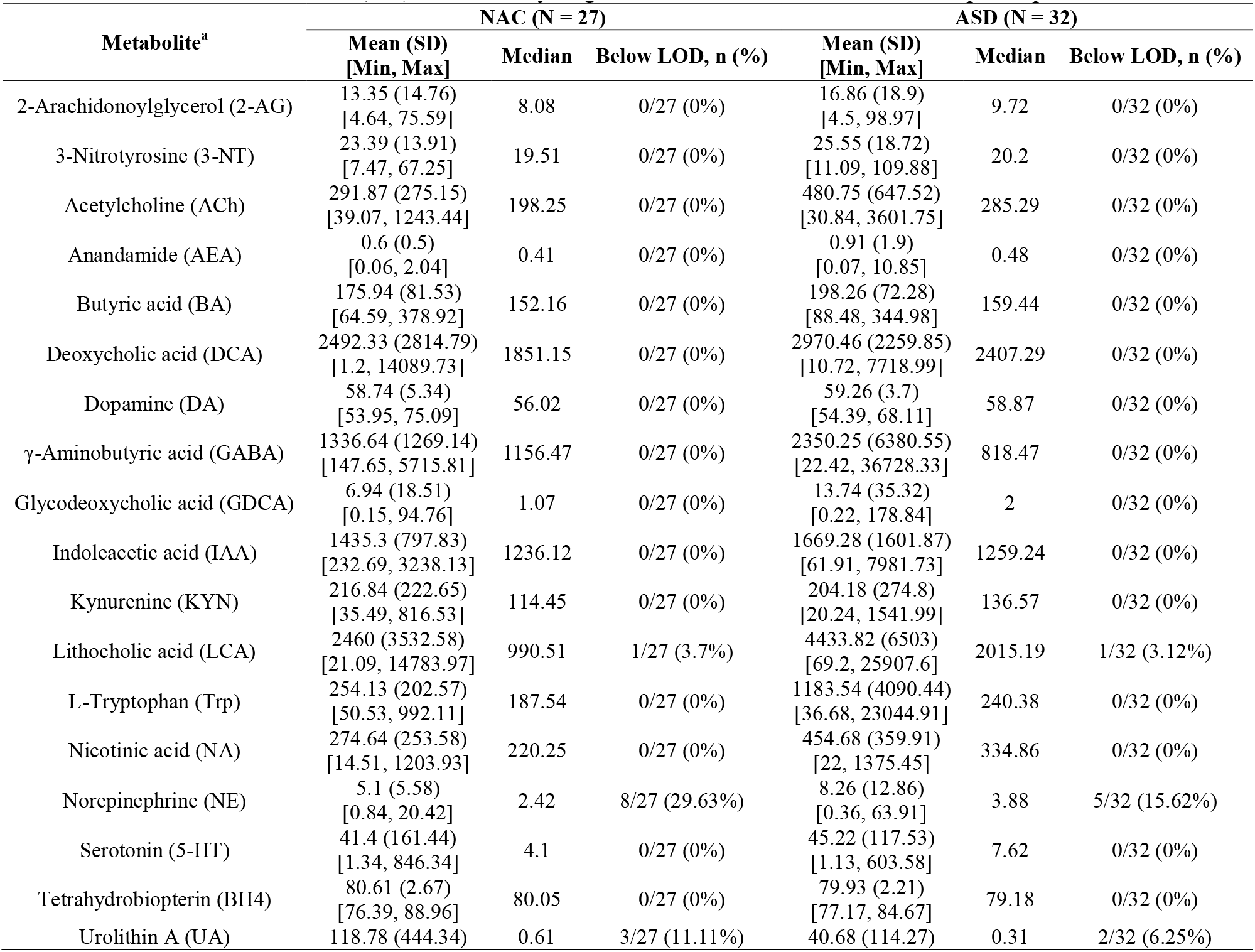
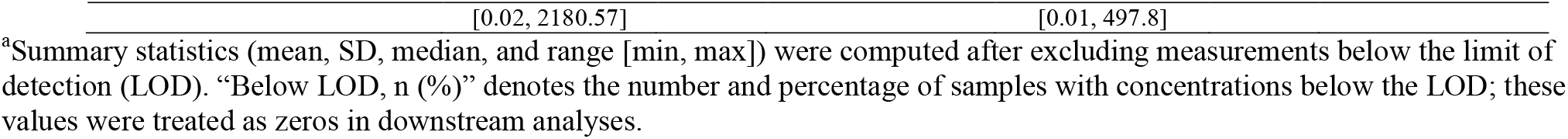
Stool metabolite concentrations (nM) measured by targeted LC-MS/MS in NAC and ASD participants.

**Figure 1.**
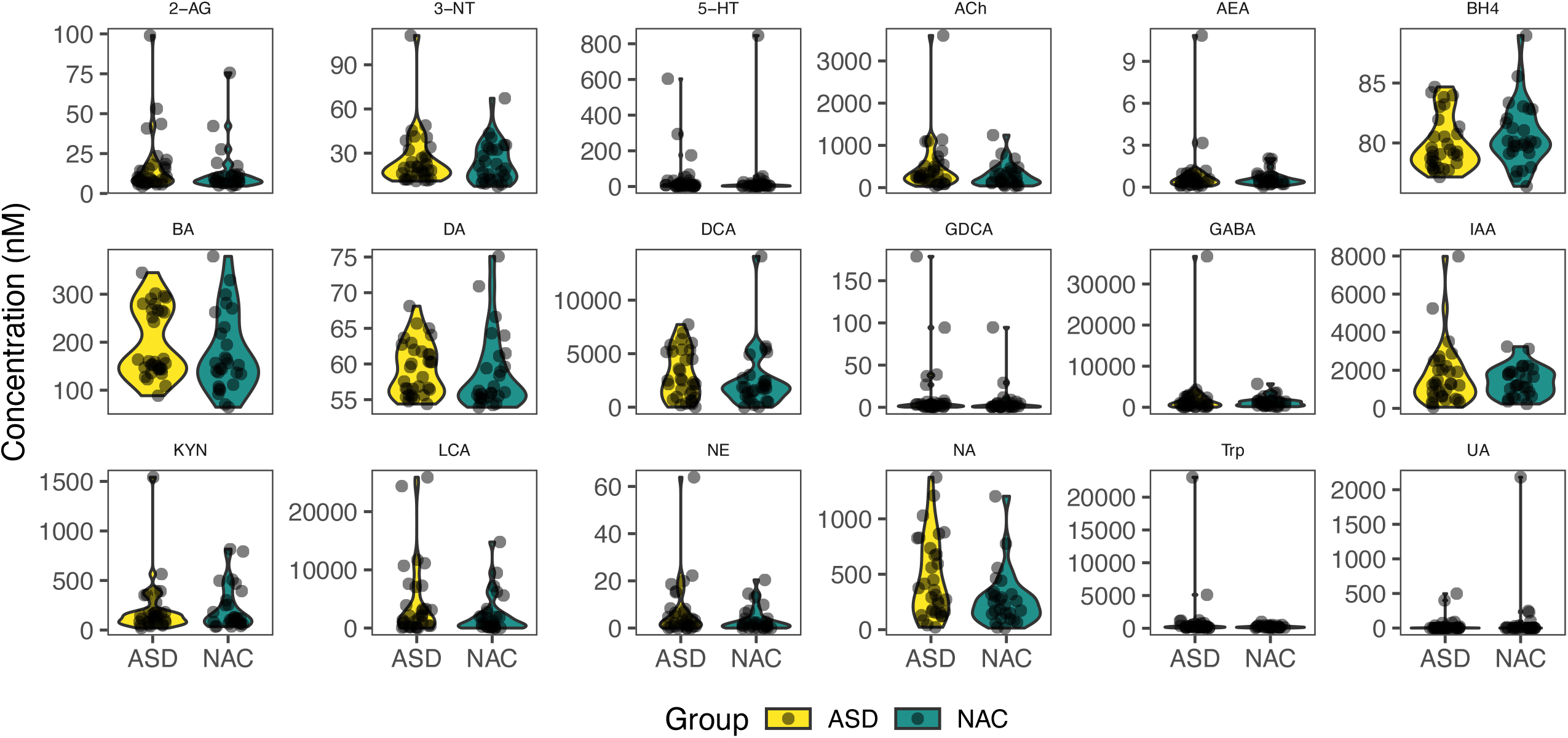
Distribution of Metabolite Concentrations in ASD and NAC Groups.

### 3.1. Groupwise Differences in Stool Metabolite Abundance and Statistical Assessment

The group-wise differences for each metabolite were assessed using log2 fold change based on normalized and log-transformed metabolite concentrations (Figure 2, Table S2). Among all metabolites, NE exhibited the largest positive log2FC (1.42), suggesting that NE levels are approximately 2.68 times higher in the ASD group relative to the NAC group, on average. The log2FC for other metabolites showed smaller or negligible differences (Table S2).

**Figure 2.**
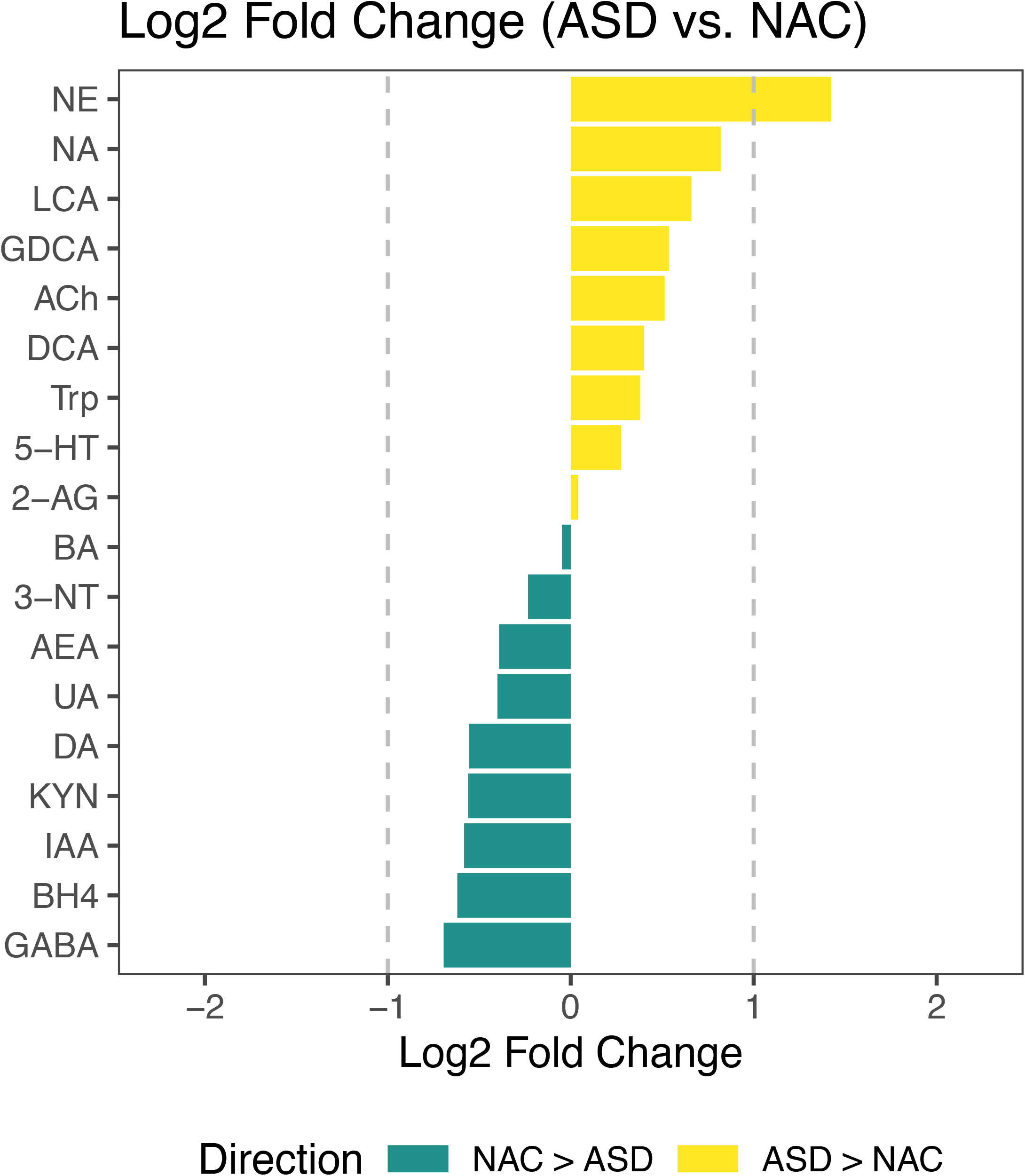
Log2 Fold Change of Metabolite Abundances Between ASD and NAC Groups. Positive values indicate higher abundance in the ASD group, while negative values indicate higher abundance in the NAC group. Dashed lines represent ±1 log2 fold change thresholds, corresponding to a two-fold difference in either direction.

Subsequently, we evaluated the statistical significance of these observed groupwise differences using linear models, with group as the predictor and age and sex as covariates. We found that DA (uncorrected p = 0.031) and BH4 (uncorrected p = 0.020) showed nominal statistical significance based on uncorrected p-values. However, these did not remain significant following adjustment for multiple comparisons using FDR (p > 0.05). The detailed model statistics are reported alongside the groupwise means and log2FC values in Table S2.

### 3.2. Discriminative Performance of Metabolite Features in Classifying ASD

To evaluate the discriminative performance of fecal metabolites in classifying ASD, we aimed to identify the most informative metabolites and construct multivariate random forest models based on these top features. As such, we first trained random forest models using each metabolite individually, evaluated them using five-fold cross-validation, and found that the top three metabolites based on the AUC with 95% CIs were BH4 (AUC = 0.683 [0.543-0.824]), GABA (AUC = 0.600 [0.452-0.747]), and KYN (AUC = 0.593 [0.445-0.740]; Table S3).

We then constructed multivariate random forest models and evaluated their performance by combining sex and age features with the same cross-validation procedure (Figure 3). Models based on age or sex alone showed limited discrimination, whereas combining the top three metabolites (BH4, KYN, and GABA) yielded an AUC of 0.737 [0.603-0.871], with an accuracy of 0.719, an F1-score of 0.670, a precision of 0.768, a recall of 0.633, and a specificity of 0.790. Adding sex to this model increased the AUC to 0.765 [0.644-0.887], whereas adding age decreased the AUC to 0.650 [0.507-0.794]. The full model incorporating age, sex, BH4, KYN, and GABA achieved an AUC of 0.688 [0.550-0.825]. A complete and detailed list of model performance metrics is provided in Table S4.

**Figure 3.**
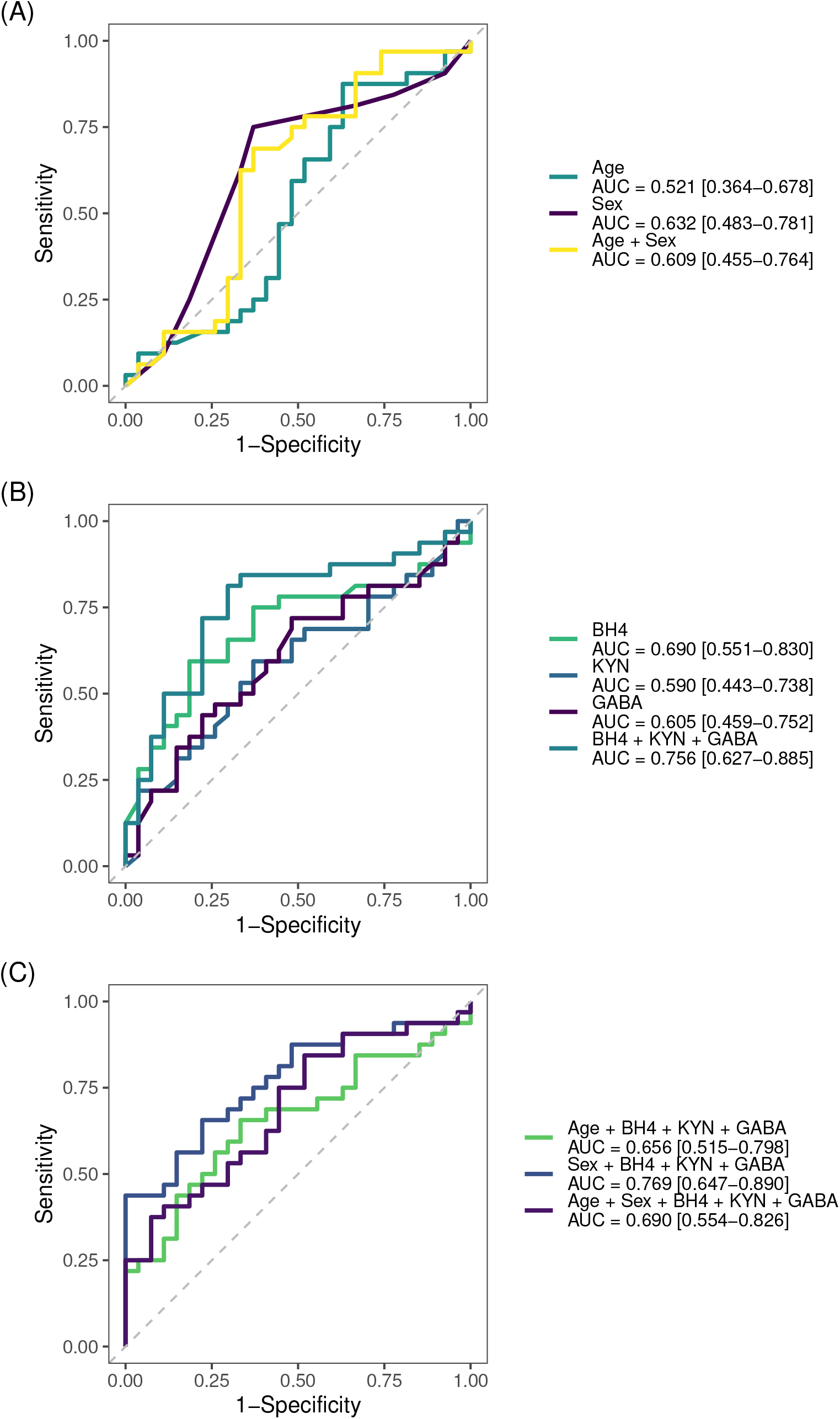
ROC curves for random forest models predicting ASD vs. NAC across different predictor sets. ROC curves showing the classification performance of random forest models based on (A) demographic variables only, (B) individual and combined metabolite markers, and (C) combined models integrating demographics with metabolites. Curves represent average performance across five-fold cross-validation. AUC values and 95% confidence intervals are shown in the legend.

### 3.3. Model Calibration Performance

To ensure that the predicted probabilities of the constructed random forest models reflect the true likelihood of outcomes, we evaluated the model calibration using the Brier score and LOESS-smoothed calibration curves (Figure 4). Brier scores quantify the mean squared difference between predicted probabilities and actual outcomes, with lower values indicating better calibration.

**Figure 4.**
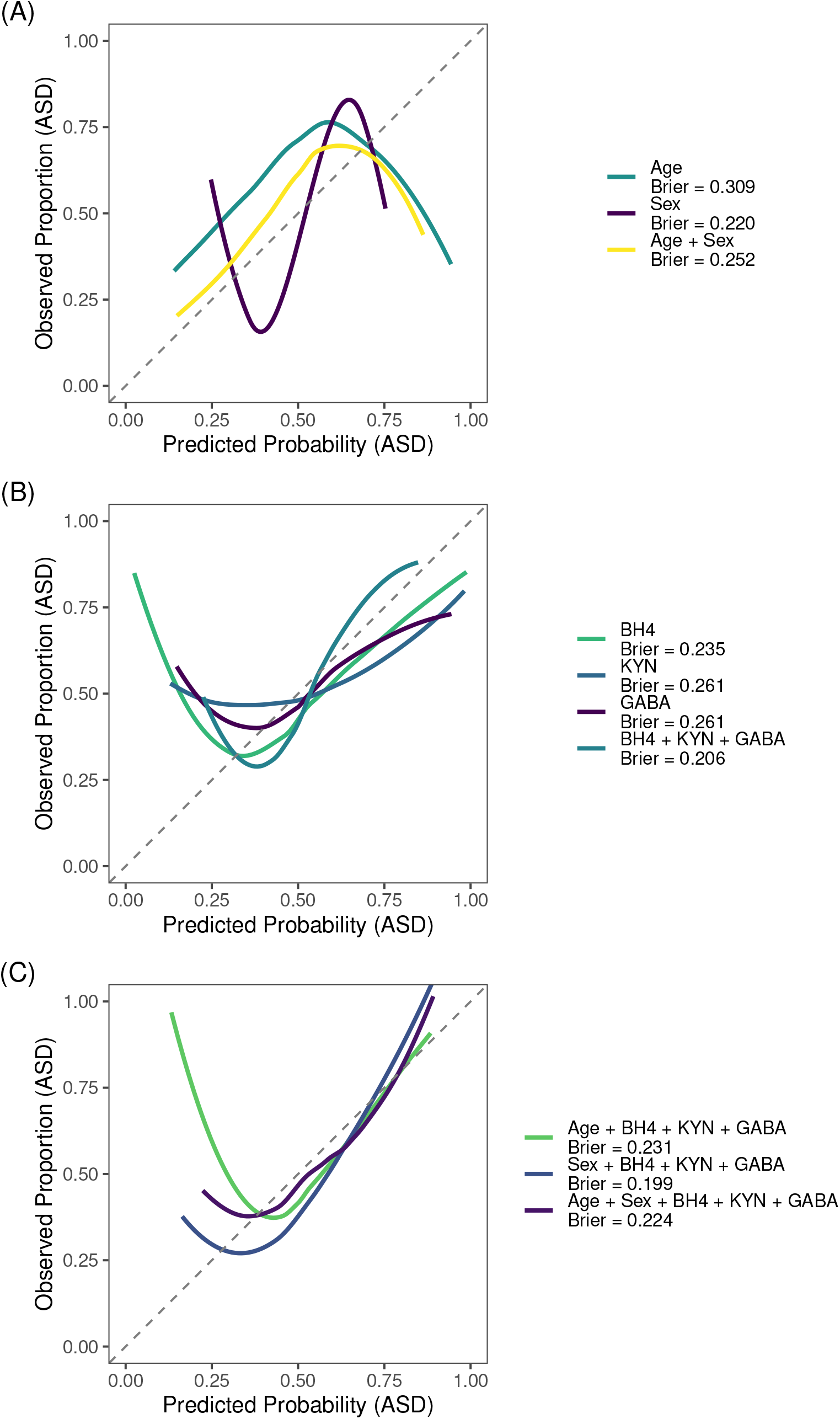
Calibration of Random Forest Models Across Predictor Sets. Calibration plots comparing predicted probabilities of ASD classification to observed outcome proportions using LOESS-smoothed curves from five-fold cross-validation for models using (A) demographic variables only, (B) individual and combined metabolite markers, and (C) combined models integrating demographics with metabolites. The gray dashed line indicates perfect calibration. Brier scores are shown in the legend.

The model combining sex with the top three metabolites (BH4, KYN, and GABA) showed the best calibration, achieving the lowest Brier score of 0.199. This was followed by the model including only the top three metabolites (Brier score = 0.206). The full model that incorporated age, sex, and the top three metabolites had a Brier score of 0.224, while the model combining age and the three metabolites had a Brier score of 0.231. Among demographic-only models, the sex-only model showed better calibration (Brier score = 0.220) than the age-only model (Brier score = 0.309), and the age + sex model had a Brier score of 0.252. Among individual metabolite models, BH4 showed better calibration (0.235) than KYN and GABA, both of which had Brier scores of 0.261. Full calibration scores for all models are reported in Table S4.

### 3.4. Within-Group Correlation Analysis of Stool Metabolites

We next examined within-group correlations among stool metabolites and age using Spearman correlation. A total of eight significant correlations were identified in the ASD group and three in the NAC group, following FDR correction applied separately within the ASD and NAC groups (Table S5). In both groups, LCA and deoxycholic acid (DCA) were strongly correlated (r = 0.85 in ASD; r = 0.78 in NAC). In the ASD group, additional significant correlations included Trp with GABA (r = 0.68), Nicotinic acid (NA, r = 0.53), anandamide (AEA; r = 0.58), and NE (r = 0.72), as well as NE with AEA (r = 0.66), NA with GABA (r = 0.63), and 3-nitrotyrosine with urolithin A (r = 0.55). In the NAC group, significant correlations were observed between Trp and AEA (r = 0.66) and between Trp and KYN (r = 0.61). The correlation matrices for each group are shown in Figure 5.

**Figure 5.**
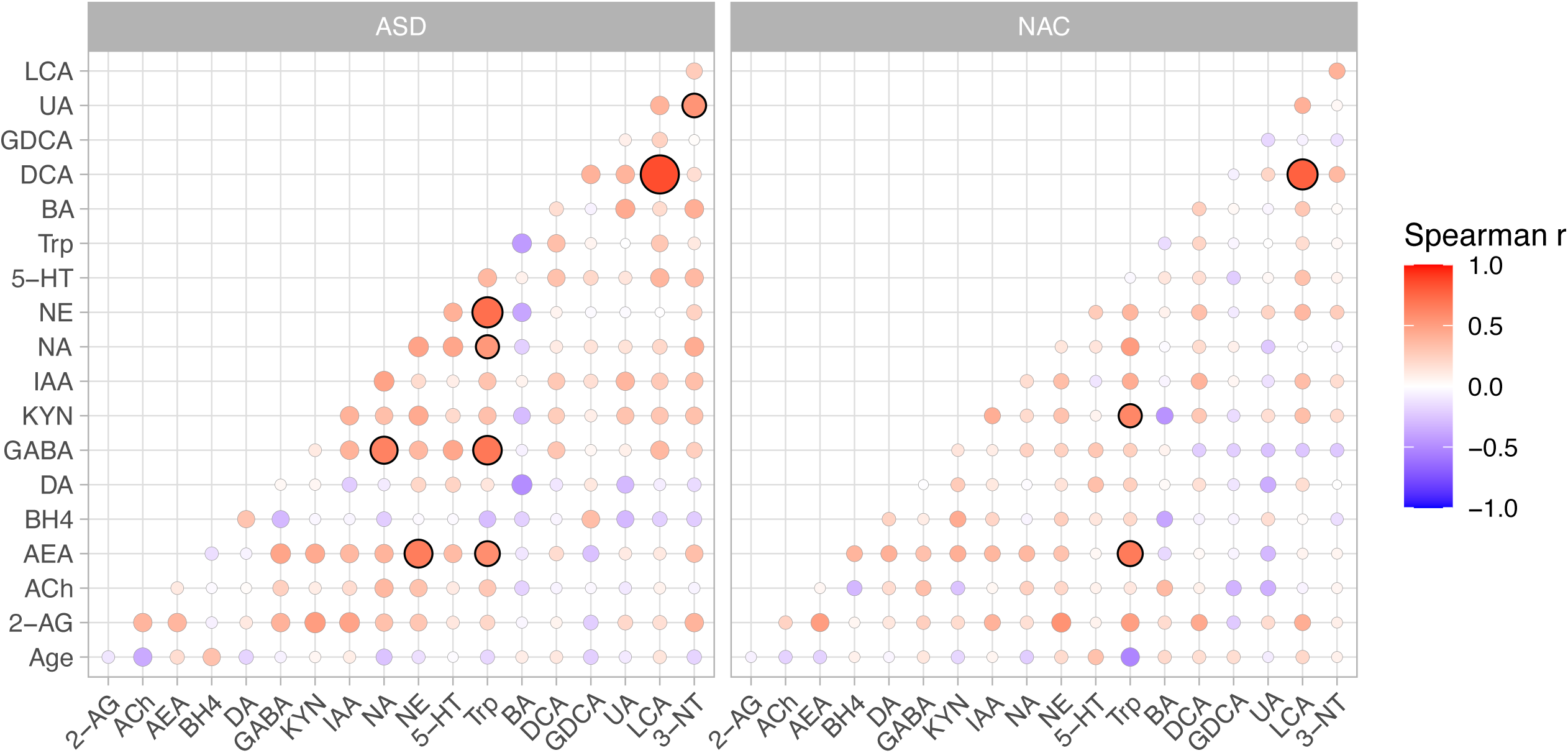
Within-group correlation matrices of stool metabolites and age in ASD and NAC. Spearman correlations are shown for ASD (left) and NAC (right). Circle color represents the correlation coefficient (r; red = positive, blue = negative), circle size indicates the strength of statistical evidence (-log_10_[p]), and significant correlations after FDR correction (p < 0.05) are highlighted with a black outline.

## 4. Discussion

### 4.1. Summary of Principal Findings

We identified NE as the largest group difference, with DA and BH4 showing nominal effects that did not survive FDR correction. Random forest analysis highlighted BH4, GABA, and KYN as the most informative metabolites, with improved performance when sex was included. Correlation analyses revealed conserved LCA-DCA coupling and altered Trp-linked associations with ASD. A small subset of metabolites (LCA, NE, UA) showed values at or below the LOD and manual inspection of MS signals confirmed these reflect true low abundance. Overall, the results indicate group-level metabolic differences and altered metabolic network structure in ASD.

### 4.2. Catecholaminergic Imbalance in ASD

The classical dopamine hypothesis of ASD attributes its core behavioral features to dysfunction within central dopaminergic circuits. (22) Dopaminergic regulation also operates peripherally, and gut microbiome-mediated processes can influence catecholamine signaling via the GBA. (23) Consistent with this, our metabolomic data showed reduced BH4 and DA and elevated NE in ASD.

The BH4 decrease is notable given its essential role in monoamine synthesis and social behavior. As a required cofactor for the biosynthesis of dopamine, serotonin, and nitric oxide, BH4 depletion can impair neurotransmitter production and synaptic signaling. (22,24) In ASD mouse models, *L. reuteri* restores social behavior by increasing BH4-pathway metabolites, and BH4 supplementation alone can reproduce this rescue. (13) Consistent with these preclinical data, pediatric studies report improvements in social and behavioral domains with BH4 supplementation. (25) Together, these observations support BH4 as a mechanistic link between gut dysbiosis, altered catecholamine synthesis, and social impairments in ASD.

The concurrent reductions in DA and BH4 align with BH4’s role as a cofactor for tyrosine hydroxylase. (26) By contrast, the elevation of stool NE diverges from this expected relationship, suggesting additional sources beyond host DA availability. One plausible explanation is partial microbial production independent of host catecholamine metabolism. (23) Several gut bacteria can synthesize NE *de novo* through enzymes analogous to dopamine β-monooxygenase. (23) NE also modulates microbial quorum sensing, reinforcing bidirectional host-microbe signaling. (23) Taken together, these findings indicate that the catecholaminergic imbalance observed in ASD likely reflects both reduced BH4-dependent DA biosynthesis and altered microbial catecholamine activity. Interventions that restore BH4 availability or modulate microbial catecholamine synthesis and signaling may help re-establish gut catecholaminergic homeostasis and mitigate downstream neurobehavioral effects. (27–29)

### 4.3. Microbiome-Linked Reorganization of Tryptophan and NAD^+^ Metabolism

To explore alterations in metabolic pathways associated with gut microbial activity, we examined within-group correlations among fecal metabolites. A robust positive correlation between LCA and DCA was observed in both groups, consistent with their shared microbial origin. (30) Beyond this conserved bile acid relationship, the most striking group differences involved Trp-linked metabolic pathways. In controls, Trp correlated strongly with KYN, consistent with coordinated activity of the hepatic KYN pathway. (31,32) However, this coupling was absent in the ASD group, suggesting altered coordination of Trp metabolism through the KYN pathway. Instead, Trp formed additional associations with NA, GABA, NE, and AEA.

The Trp-NA association is noteworthy because both metabolites are integral to nicotinamide adenine dinucleotide (NAD^+^) biosynthesis. NAD^+^ can be produced *de novo* from Trp through the KYN pathway or from NA via the Preiss-Handler deamidated route. (33) Microbial nicotinamidase (PncA) can bypass host constraints by converting nicotinamide (NAM) to NA, feeding into the Preiss-Handler pathway. (34) Since mammalian cells lack the ability to deamidate NAM, this microbial reaction creates a unique host-microbe interface in NAD^+^ metabolism.

This microbial contribution to host NAD^+^ metabolism provides a plausible explanation for the altered Trp-NA coupling observed in ASD. This correlation may reflect enhanced or dysregulated microbial participation in NAD^+^ precursor cycling, potentially as a compensatory response to maintain NAD^+^ levels under conditions of disrupted Trp-KYN metabolism or oxidative imbalance. Alternatively, shifts in the abundance or activity of PncA-expressing taxa could drive excessive deamidation, favoring NA-dependent NAD^+^ biosynthesis over canonical host routes. Collectively, these results highlight a host-microbe metabolic axis linking NAD^+^ homeostasis to gut microbial activity in ASD.

### 4.4. Discriminative Value of Stool Metabolites

Motivated by the catecholaminergic and Trp-linked disruptions described above and the feasibility of stool as a pediatric, noninvasive sample, we assessed whether stool metabolites carry discriminative signal for ASD. Prior work suggests that gut-microbiome features can aid ASD screening and subtype stratification. (35–38) Building on this foundation, we extend from taxa-centric to metabolite-centric profiling, offering a complementary readout that may be informative when behavioral assessments are least reliable. Among the profiled metabolites, BH4, GABA, and KYN were the most informative and map to non-redundant biology, which likely explains their combined gain in discrimination. BH4 captures cofactor availability for monoamine synthesis and aligns with the catecholaminergic findings above (24) ; GABA reflects inhibitory neurotransmission that can be produced and modulated by gut microbes (39) ; and KYN marks immune-regulated Trp catabolism, consistent with the altered Trp-linked correlations we observed (31,32) . Notably, each single-metabolite model showed only modest discrimination (AUC < 0.7), indicating that no single compound is sufficient; the improvement arises from non-redundant biology across these pathways. The additional boost when sex is included is directionally consistent with known sex differences in ASD prevalence and gut-brain physiology. (40) By contrast, age reduced discrimination, likely due to the cohort’s uneven age structure, which introduces non-monotonic patterns and adds variance without stable diagnostic signal. Overall, the multimetabolite panel provided moderate discrimination with acceptable calibration, supporting its value as a mechanistically coherent, hypothesis-generating signature rather than a diagnostic test.

### 4.5. Study Limitations and Future Directions

Several limitations of this study should be acknowledged. First, the sample size was modest, which limits statistical power to detect small-to-moderate effects. While multiple metabolites showed differences between groups, none survived false discovery rate correction, and the findings should therefore be considered exploratory. Second, the cross-sectional design precludes causal inference or assessment of developmental trajectories, which require longitudinal follow-up. Third, potential confounders such as diet, gastrointestinal symptoms, medication use, and underlying microbiome composition were not directly measured or controlled and may influence the stool metabolome. (41–43) Pre-analytical and analytical factors (collection timing, storage, matrix effects, batch variation) may also introduce variability. Although below-LOD values were confirmed to reflect true low-abundance signals, they were excluded from descriptive summaries but retained as zeros in downstream analyses, which may attenuate subtle differences. The cohort’s age structure and sex imbalance may confound associations, and the machine-learning models require external validation. In addition, participants with ASD were recruited via clinician referral and specialty clinical programs, whereas non-ASD controls were recruited from community sources. This convenience sampling strategy may introduce selection bias and limit the representativeness of both groups relative to the broader pediatric population. Differences in healthcare-seeking behavior, socioeconomic factors, or unmeasured environmental exposures between recruitment sources may further influence observed metabolic patterns despite statistical adjustment for age and sex.

Future studies should incorporate larger, independent cohorts and longitudinal sampling to assess developmental trajectories. Integrative multi-omics approaches may clarify mechanisms linking metabolic alterations to ASD, and translational work is needed to evaluate stool metabolites as adjunctive biomarkers and to test targeted metabolic or microbiome-based interventions. (25,27,44)

Despite these limitations, this study advances efforts to characterize the stool metabolome in ASD. We identify network-level alterations involving Trp, pterin, and catecholamine pathways and show that stool metabolites capture clinically relevant variation. These preliminary findings provide a foundation for clarifying gut-derived metabolic contributions to neurodevelopment and for evaluating their biomarker potential.

## Acknowledgements

We sincerely thank all participants and their parents for their invaluable contributions to this project. We are grateful to Dr. Bruce Rosen for his support throughout the study and to Dr. Chongzhao Ran for laboratory assistance. We thank Dr. Spyros Nikas for providing anandamide and 2-arachidonoylglycerol used in the preparation of calibration and quality control samples.

This research was funded by Boston Children’s Hospital (Grant Agreement 92436) and Massachusetts General Hospital (233263), with Dr. Xue-Jun Kong as principal investigator. The funders had no role in study design, data collection, analysis, manuscript preparation, or the decision to submit the work for publication.

## Author Contributions

Conceptualization: X.-J.K., K.L.; methodology: X.-J.K., K.L., J.J.G., H.L.; software: K.L.; validation: K.L.; formal analysis: K.L.; investigation: X.-J.K., J.J.G., K.L., H.L., S.Z., M.X., J.Z., J.C., W.X., A.X.; resources: X.-J.K.; J.J.G.; data curation: K.L., H.L.; writing—original draft preparation: K.L.; writing—review and editing: K.L., H.L., S.Z., M.X., J.Z., J.C., W.X., A.X., A.M., J.J.G., X.-J.K.; visualization: K.L.; supervision: X.-J.K., J.J.G.; project administration: X.-J.K., J.J.G.; funding acquisition: X.-J.K.; all authors have read and agreed to the published version of the manuscript.

## Ethics Approval

The study was conducted in accordance with the Declaration of Helsinki and approved by the Institutional Review Board of Massachusetts General Hospital (protocol 2022P001749, approval date: August 8, 2022).

## Informed Consent

Written informed consent was obtained from all subjects involved in the study and assent was obtained when appropriate.

## Legends for Figures

**Figure S1.** Age Distribution of Study Participants by Diagnostic Group.

## References

1. American-Psychiatric-Association (2013): Diagnostic and Statistical Manual of Mental Disorders: DSM-5, 5th edition, vol. 5. Washington, DC: American psychiatric association.

2. Luo Y, Eran A, Palmer N, Avillach P, Levy-Moonshine A, Szolovits P, Kohane IS (2020): A multidimensional precision medicine approach identifies an autism subtype characterized by dyslipidemia. Nat Med 26: 1375–1379.

3. Feczko E, Balba NM, Miranda-Dominguez O, Cordova M, Karalunas SL, Irwin L, et al. (2018): Subtyping cognitive profiles in Autism Spectrum Disorder using a Functional Random Forest algorithm. NeuroImage 172: 674–688.

4. Mosconi MW, Stevens CJ, Unruh KE, Shafer R, Elison JT (2023): Endophenotype trait domains for advancing gene discovery in autism spectrum disorder. J Neurodev Disord 15: 41.

5. Lord C, Rutter M, Couteur AL (1994): Autism Diagnostic Interview-Revised: A revised version of a diagnostic interview for caregivers of individuals with possible pervasive developmental disorders. J Autism Dev Disord 24: 659–685.

6. Lord C, Risi S, Lambrecht L, Jr EHC, Leventhal BL, DiLavore PC, et al. (2000): The Autism Diagnostic Observation Schedule—Generic: A standard measure of social and communication deficits associated with the spectrum of autism. Journal of autism and developmental disorders 30: 205–223.

7. Parellada M, Andreu-Bernabeu Á, Burdeus M, Cáceres ASJ, Urbiola E, Carpenter LL, et al. (2023): In Search of Biomarkers to Guide Interventions in Autism Spectrum Disorder: A Systematic Review. Am J Psychiatry 180: 23–40.

8. Carabotti M, Scirocco A, Maselli MA, Severi C (2014): The gut-brain axis: interactions between enteric microbiota, central and enteric nervous systems. Ann Gastroenterol 28: 203–209.

9. Petrut S-M, Bragaru AM, Munteanu AE, Moldovan A-D, Moldovan C-A, Rusu E (2025): Gut over Mind: Exploring the Powerful Gut–Brain Axis. Nutrients 17: 842.

10. Zheng L, Jiao Y, Zhong H, Tan Y, Yin Y, Liu Y, et al. (2024): Human-derived fecal microbiota transplantation alleviates social deficits of the BTBR mouse model of autism through a potential mechanism involving vitamin B6 metabolism. mSystems 9: e00257–24.

11. Kang D-W, Adams JB, Vargason T, Santiago M, Hahn J, Krajmalnik-Brown R (2020): Distinct Fecal and Plasma Metabolites in Children with Autism Spectrum Disorders and Their Modulation after Microbiota Transfer Therapy. mSphere 5: 10.1128/msphere.00314-20.

12. Kang D-W, Adams JB, Gregory AC, Borody T, Chittick L, Fasano A, et al. (2017): Microbiota Transfer Therapy alters gut ecosystem and improves gastrointestinal and autism symptoms: an open-label study. Microbiome 5: 10.

13. Buffington SA, Dooling SW, Sgritta M, Noecker C, Murillo OD, Felice DF, et al. (2021): Dissecting the contribution of host genetics and the microbiome in complex behaviors. Cell 184: 1740-1756.e16.

14. Dooling SW, Sgritta M, Wang I-C, Duque ALRF, Costa-Mattioli M (2022): The Effect of Limosilactobacillus reuteri on Social Behavior Is Independent of the Adaptive Immune System. mSystems 7: e00358–22.

15. Gevi F, Belardo A, Zolla L (2020): A metabolomics approach to investigate urine levels of neurotransmitters and related metabolites in autistic children. Biochim Biophys Acta (BBA) - Mol Basis Dis 1866: 165859.

16. Wang D, Jiang Y, Jiang J, Pan Y, Yang Y, Fang X, et al. (2025): Gut microbial GABA imbalance emerges as a metabolic signature in mild autism spectrum disorder linked to overrepresented Escherichia. Cell Rep Med 6: 101919.

17. Kałużna-Czaplińska J, Gątarek P, Chirumbolo S, Chartrand MS, Bjørklund G (2019): How important is tryptophan in human health? Crit Rev Food Sci Nutr 59: 72–88.

18. Luna RA, Oezguen N, Balderas M, Venkatachalam A, Runge JK, Versalovic J, et al. (2017): Distinct Microbiome-Neuroimmune Signatures Correlate With Functional Abdominal Pain in Children With Autism Spectrum Disorder. Cell Mol Gastroenterol Hepatol 3: 218–230.

19. Hu T, Dong Y, He C, Zhao M, He Q (2020): The Gut Microbiota and Oxidative Stress in Autism Spectrum Disorders (ASD). Oxidative Med Cell Longev 2020: 8396708.

20. Vuong HE, Hsiao EY (2017): Emerging Roles for the Gut Microbiome in Autism Spectrum Disorder. Biol Psychiatry 81: 411–423.

21. Pang Z, Lu Y, Zhou G, Hui F, Xu L, Viau C, et al. (2024): MetaboAnalyst 6.0: towards a unified platform for metabolomics data processing, analysis and interpretation. Nucleic Acids Res 52: W398–W406.

22. Pavăl D (2017): A Dopamine Hypothesis of Autism Spectrum Disorder. Dev Neurosci 39: 355–360.

23. Miri S, Yeo J, Abubaker S, Hammami R (2023): Neuromicrobiology, an emerging neurometabolic facet of the gut microbiome? Front Microbiol 14: 1098412.

24. Werner ER, Blau N, Thöny B (2011): Tetrahydrobiopterin: biochemistry and pathophysiology. Biochem J 438: 397–414.

25. Klaiman C, Huffman L, Masaki L, Elliott GR (2013): Tetrahydrobiopterin as a Treatment for Autism Spectrum Disorders: A Double-Blind, Placebo-Controlled Trial. J Child Adolesc Psychopharmacol 23: 320–328.

26. Fanet H, Capuron L, Castanon N, Calon F, Vancassel S (2021): Tetrahydrobioterin (BH4) Pathway: From Metabolism to Neuropsychiatry. Curr Neuropharmacol 19: 591–609.

27. Frye RE, DeLaTorre R, Taylor HB, Slattery J, Melnyk S, Chowdhury N, James SJ (2013): Metabolic effects of sapropterin treatment in autism spectrum disorder: a preliminary study. Transl Psychiatry 3: e237–e237.

28. Clarke MB, Hughes DT, Zhu C, Boedeker EC, Sperandio V (2006): The QseC sensor kinase: A bacterial adrenergic receptor. Proc Natl Acad Sci 103: 10420–10425.

29. Liu L, Huh JR, Shah K (2022): Microbiota and the gut-brain-axis: Implications for new therapeutic design in the CNS. eBioMedicine 77: 103908.

30. Kiriyama Y, Nochi H (2021): Physiological Role of Bile Acids Modified by the Gut Microbiome. Microorganisms 10: 68.

31. Badawy AA-B (2017): Kynurenine Pathway of Tryptophan Metabolism: Regulatory and Functional Aspects. Int J Tryptophan Res 10: 1178646917691938.

32. Dehhaghi M, Panahi HKS, Guillemin GJ (2019): Microorganisms, Tryptophan Metabolism, and Kynurenine Pathway: A Complex Interconnected Loop Influencing Human Health Status. Int J Tryptophan Res 12: 1178646919852996.

33. Xie N, Zhang L, Gao W, Huang C, Huber PE, Zhou X, et al. (2020): NAD+ metabolism: pathophysiologic mechanisms and therapeutic potential. Signal Transduct Target Ther 5: 227.

34. Shats I, Williams JG, Liu J, Makarov MV, Wu X, Lih FB, et al. (2020): Bacteria Boost Mammalian Host NAD Metabolism by Engaging the Deamidated Biosynthesis Pathway. Cell Metab 31: 564-579.e7.

35. Kong X, Liu J, Cetinbas M, Sadreyev R, Koh M, Huang H, et al. (2019): New and Preliminary Evidence on Altered Oral and Gut Microbiota in Individuals with Autism Spectrum Disorder (ASD): Implications for ASD Diagnosis and Subtyping Based on Microbial Biomarkers. Nutrients 11: 2128.

36. Wan Y, Wong OWH, Tun HM, Su Q, Xu Z, Tang W, et al. (2024): Fecal microbial marker panel for aiding diagnosis of autism spectrum disorders. Gut Microbes 16: 2418984.

37. Kong X, Liu J, Liu K, Koh M, Tian R, Hobbie C, et al. (2021): Altered Autonomic Functions and Gut Microbiome in Individuals with Autism Spectrum Disorder (ASD): Implications for Assisting ASD Screening and Diagnosis. J Autism Dev Disord 51: 144–157.

38. Su Q, Wong OWH, Lu W, Wan Y, Zhang L, Xu W, et al. (2024): Multikingdom and functional gut microbiota markers for autism spectrum disorder. Nat Microbiol 9: 2344–2355.

39. Braga JD, Thongngam M, Kumrungsee T (2024): Gamma-aminobutyric acid as a potential postbiotic mediator in the gut–brain axis. npj Sci Food 8: 16.

40. Kopec AM, Fiorentino MR, Bilbo SD (2018): Gut-immune-brain dysfunction in Autism: Importance of sex. Brain Res 1693: 214–217.

41. Zierer J, Jackson MA, Kastenmüller G, Mangino M, Long T, Telenti A, et al. (2018): The fecal metabolome as a functional readout of the gut microbiome. Nat Genet 50: 790–795.

42. Shinn LM, Mansharamani A, Baer DJ, Novotny JA, Charron CS, Khan NA, et al. (2022): Fecal Metabolites as Biomarkers for Predicting Food Intake by Healthy Adults. J Nutr 152: 2956–2965.

43. Needham BD, Adame MD, Serena G, Rose DR, Preston GM, Conrad MC, et al. (2021): Plasma and Fecal Metabolite Profiles in Autism Spectrum Disorder. Biol Psychiatry 89: 451–462.

44. Filho CC, Melfior L, Ramos SL, Pizi MSO, Taruhn LF, Muller ME, et al. (2025): Tetrahydrobiopterin and Autism Spectrum Disorder: A Systematic Review of a Promising Therapeutic Pathway. Brain Sci 15: 151.

